# *PgaR* is a positive regulator of the *pgaABCD* biosynthetic operon in *Klebsiella pneumoniae*

**DOI:** 10.64898/2026.03.13.711557

**Authors:** Jonathan Bradshaw, Julia Sanchez-Garrido, Sophia David, Mariagrazia Pizza, Immaculada Margarit Ros, Maria Rosaria Romano, Joshua L.C. Wong, Gad Frankel

## Abstract

The biosynthetic locus encoding the exopolysaccharide poly-N-acetyl-glucosamine (PNAG) is widely conserved across bacteria, including the WHO critical-priority pathogen *Klebsiella pneumoniae* (Kp). In Kp, PNAG synthesis is mediated by the *pgaABCD* operon, yet its lineage-specific regulation remains incompletely defined. Using a comparative genomics approach to interrogate the *pgaABCD* locus across the high-risk clonal Kp complex 258 (CC258) lineage, we identified a previously uncharacterised positive transcriptional regulator located immediately upstream of *pgaA*, which we designate *pgaR*. Phylogenetic analysis revealed recurrent evolutionary events affecting this regulatory region, including repeated deletion or truncation of *pgaR* and a G>A substitution upstream of the *pgaR* start codon. Functional characterisation demonstrated that loss of *pgaR* abolishes *pgaABCD* expression and PNAG production, whereas the upstream G>A substitution drives PNAG hyper-production. In vitro, Kp produce extensive extracellular PNAG networks under static growth conditions, consistent with a role in biofilm architecture. Despite this, PNAG expression was dispensable in murine pneumonia and peritonitis models, while PNAG hyper-production significantly attenuated virulence and disease severity, indicating a fitness cost associated with sustained overexpression. Collectively, we discovered PgaR as a novel gene regulator of the *pgaABCD* operon. We show a previously unrecognised lineage-specific layer of PNAG regulation in Kp and demonstrate that opposing PNAG phenotypes: loss and hyper-production, have independently and repeatedly emerged among clinical CC258 isolates, highlighting dynamic selection acting on biofilm-associated traits in this high-risk pathogen.

**Importance:** The exopolysaccharide poly-N-acetyl-glucosamine (PNAG) is widely conserved in bacteria, including the WHO critical-priority pathogen *Klebsiella pneumoniae*. However, how PNAG production is regulated in high-risk lineages has remained unclear. Here, we identify PgaR as a previously unrecognised positive regulator of the *pgaABCD* operon in clonal complex 258, a globally disseminated and drug-resistant lineage. We show that natural genetic variation within this regulatory region leads to strikingly different PNAG phenotypes: complete loss of production or hyper-production. While PNAG contributes to extracellular matrix formation in vitro, it is dispensable for virulence in murine infection models, and sustained overproduction imposes a fitness cost. The repeated and independent emergence of both loss- and gain-of-function variants among clinical isolates reveals dynamic evolutionary pressures acting on biofilm-associated traits. These findings uncover a lineage-specific layer of PNAG regulation and highlight how modulation of surface polysaccharide expression shapes pathogen fitness and adaptation.

## Introduction

Bacterial surface polysaccharides often contribute to virulence and represent attractive therapeutic targets. Poly-N-Acetyl-Glucosamine (PNAG) is a high molecular weight bacterial polysaccharide consisting of repeat units of N-Acetyl-Glucosamine (GlcNAc) joined via a β(1–6) glycosidic linkage (1,2). Unlike other bacterial surface molecules, which are species specific (e.g. lipopolysaccharide O-antigen), PNAG is conserved across bacteria, commensals and pathogens alike, and has been detected in most members of the World Health Organisation’s (WHO) critical and high priority pathogen list (3–6).

PNAG production in the encapsulated Gram-negative pathogen *Klebsiella pneumoniae* (Kp) has not been thoroughly investigated. Kp is a common constituent of the human microbiome, where it colonises the mucosal surfaces of the oropharynx and gastrointestinal (GI) tract (7). Classical Kp (cKp) cause infections in immunocompromised patients, commonly bacterial pneumonia and bloodstream infections, and are associated with broad antimicrobial resistance (8). Of particular concern are infections caused by cKp resistant to carbapenems and third generation cephalosporins, which are difficult to treat and thus, have been assigned critical priority status by the WHO (9). To date, strains of clonal complex (CC) 258, a globally distributed high-risk lineage that includes sequence type (ST) 11, 258 and 512, have been a major driver of carbapenem resistance spread in cKp (10). By contrast, hypervirulent Kp (hvKp) cause community-acquired infections primarily in healthy individuals and are associated with overproduction of the capsular polysaccharide (CPS) and carriage of large virulence plasmids (11,12). Although cKp and hvKp strains have historically been viewed as distinct pathotypes, recent convergence events, exemplified by ST11 isolates in China, have resulted in hvKp strains that exhibit multi-drug resistance (13,14).

In Gram-negative bacteria, the biosynthetic PNAG genes are organised in a four gene *pgaABCD* operon (15). In Gram-positive bacteria the *icaADBC* structural biosynthetic PNAG genes lie downstream of a negative transcriptional regulator *icaR,* which binds to sites upstream of *icaA* and inhibits *icaADBC* transcription (16,17). While the *pgaABCD* locus is the target of global transcriptional and translational regulators, no *icaR* homolog, or any PNAG-specific regulator has been identified in Gram-negative bacteria (18).

During PNAG synthesis in Gram-negative bacteria, an inner membrane complex forms between the glycosyltransferase PgaC and the accessory protein PgaD, which polymerises UDP-GlcNAc to produce a fully acetylated PNAG polymer (19,20). This polymer extends into the periplasm, where the deacetylase PgaB removes approximately 10% of the acetyl residues to facilitate export through the outer membrane porin PgaA (21). Although the *pgaABCD* locus is prevalent across Kp genomes, how conserved the gene sequences are across Kp strains has not been investigated (22). Furthermore, while immunofluorescence staining has been successful in visualising PNAG production on many bacteria, attempts to visualise PNAG production on Kp have yielded inconclusive results (23,24).

PNAG is also a known virulence factor in diverse bacterial pathogens, including Kp. For example, deletion of *pgaC* results in reduced colonisation of the GI tract following oral inoculation and reduced lethality following intraperitoneal (IP) challenge (25). Furthermore, prophylactic treatment with the PNAG-specific monoclonal antibody (mAb) F598 or polyclonal sera raised to PNAG antigen, protected mice against IP Kp challenge (5). However, the contribution of PNAG to Kp virulence at other infection sites remains unexplored. This includes pneumonia and bloodstream infections where Kp is a leading cause of mortality (26).

By analysing a large collection of Kp CC258 genomes, we found the intergenic region (IR) upstream of the PNAG biosynthetic locus to be a mutational hotspot, and within this region, identified a novel positive regulator of PNAG production (*pgaR*). Across this lineage, we identify recurrent mutations in the *pgaR* gene sequence and upstream region that result in either loss of PNAG, or a PNAG hyper-producing phenotype. Unlike other Kp surface polysaccharides which coat the bacterial surface (e.g. capsule and LPS), we show that PNAG is produced as large extracellular networks across diverse Kp strains, and that *pgaR* is required for this production. While loss of PNAG does not affect Kp virulence during *in vivo* models of Kp pneumonia and peritonitis, we show that PNAG hyper-production attenuates virulence and disease in these infection contexts. Overall, we describe a novel layer of PNAG regulation specific to Kp and demonstrates that opposing PNAG phenotypes, via mutations in *pgaR* and its upstream region, repeatedly emerge among clinical isolates.

## Results

### Kp produces extracellular PNAG networks

Current approaches to detect PNAG are limited to immunofluorescent (IF) staining, which is qualitative and prone to imaging bias. For this reason, we developed a quantitative readout of PNAG production using a combination of standardised, quantitative IF staining and enzyme-linked immunosorbent assay (ELISA).

We started using the wild-type (WT) hvKp strain ICC8001, a derivative of the prototype hvKP strain ATCC43816, which encodes an intact *pgaABCD* locus. We also generated an open reading frame (ORF) deletion mutant in the PNAG biosynthetic genes (Δ*pgaBCD*). We grew the WT and Δ*pgaBCD* strains statically on glass slides, conditions known to induce PNAG production in other bacteria, and stained with the PNAG-specific mAb F598 (1,3). We then imaged at nine standardised fields distributed evenly across the well. This revealed that the WT strain produced detectable PNAG in each field, although the quantity of PNAG varied between fields (Fig. 1A and S1A). By contrast, no PNAG was detected in Δ*pgaBCD-*coated wells (Fig. 1A and S1B). Across WT-coated wells, PNAG appeared as diffuse extracellular networks often forming long fibres between adherent cells (Fig. 1A).

**Figure 1.**
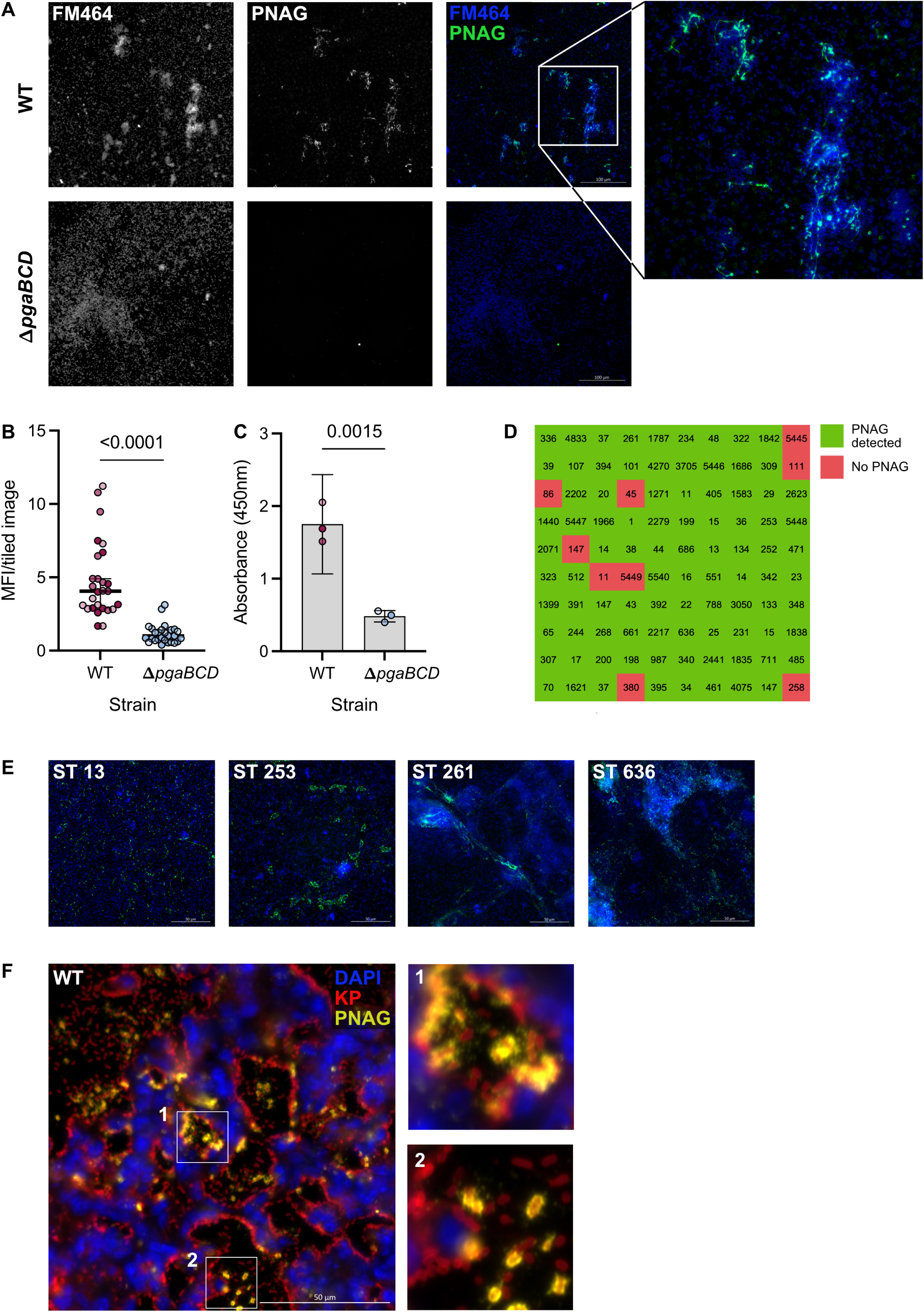
Kp produces extracellular PNAG networks. **(A)** IF staining of PNAG in WT and Δ*pgaBCD* grown statically O/N on glass slides in LB. Kp are visualised with the membrane dye FM464 (blue) and PNAG using the mAb F598 (green), with a zoomed example of PNAG extracellular network formation. Images shown are individual tiles highlighted in red in Fig. S1A and B. **(B)** Quantification of fluorescence in the PNAG channel from each of nine well locations across 3 biological replicates. MFI was calculated using FIJI image analysis software, using the default parameters for thresholding. Data is presented as median with 95% confidence intervals, and significance was assessed using a Mann-Whitney test. Individual points are coloured in different shades to represent biological repeats. **(C)** ELISA absorbance readings from wells coated with WT and Δ*pgaBCD*, probed with 500 ng/mL mAb F598. Data are presented as mean with confidence intervals, and significance was assessed using unpaired t-test. Data points are shaded to represent biological replicates. **(D)** Heat map of 100 Kp clinical isolates, identified by ST, showing the presence (green) or absence (red) of PNAG during static growth *in vitro.* Assessment of PNAG production was carried out using IF staining across two biological replicates. **(E)** Representative images of PNAG extracellular network formation across MRSN isolates, with their respective ST stated. Isolates were stained with FM464 (blue) and mAb F598 (green). A single representative image from 2 independent experiments is shown. **(F)** IF staining of Kp and PNAG in a representative WT-infected lung at 72 hpi. Sections are stained with DAPI (blue), Kp antisera (red) and mAb F598 (yellow). Boxes 1 and 2 show zoomed regions of PNAG extracellular network formation on alveolar structures, and cell-associated PNAG production, respectively.

For quantification, we measured PNAG-specific fluorescence and quantified total PNAG by conducting ELISA directly in bacterial coated wells. Consistent with IF staining, median fluorescent intensity (MFI) of WT-coated wells was significantly higher than Δ*pgaBCD-*coated wells (Fig. 1B). Furthermore, probing wells coated with WT and Δ*pgaBCD* with 500 ng/mL F598 resulted in absorbances of 1.75 and 0.5 by ELISA, respectively (Fig. 1C). This confirmed that Kp produces extracellular PNAG networks, and that deletion of the *pgaBCD* genes abolishes PNAG production.

We next investigated if the PNAG extracellular network phenotype is conserved across Kp strains. For this, we visualised PNAG production across a panel of 100 diverse Kp clinical isolates, the MRSN collection (27). After static growth on glass slides, we observed PNAG on 91 out of 100 Kp strains, spanning diverse STs (Fig. 1D). Importantly, in each case where PNAG was detected, it presented as extracellular networks, similar to WT ICC8001 (Fig. 1E). While we did observe variability in PNAG architecture across strains, we did not encounter a cell-associated phenotype (Fig. 1E). Taken together, this confirms that PNAG extracellular network formation is a conserved phenotype across diverse Kp strains.

### PNAG is produced in vivo during Kp lung infection

We next assessed if PNAG is produced by Kp during severe lung infection. To this end, we infected CD-1 mice with WT or Δ*pgaBCD* via intratracheal (IT) intubation and visualised PNAG by IF of lung sections at 72 h post infection (hpi). Staining of the WT-infected section revealed strong DAPI staining and a concomitant reduction in podoplanin staining in the dorsal portion of the lung. This is reflective of immune cell recruitment and lung epithelia disruption, respectively (Fig. S2A). We also observed strong Kp staining in this region, and interestingly, were able to detect PNAG across the breadth of the Kp-rich dorsal portion of the lung (Fig. S2A). In comparable regions of Δ*pgaBCD*-infected lung we observed the same features with absent PNAG staining (Fig. S2B). Therefore, Kp produces PNAG during *in vivo* lung infection in CD-1 mice. Upon closer inspection of PNAG in the infected lung, we observed a mixture of PNAG extracellular networks and cell-associated PNAG (Fig. 1F). While PNAG extracellular networks appeared to coat alveolar wall structures, Kp producing cell-associated PNAG appeared predominantly in the alveolar spaces (Fig. 1F).

### The intergenic *csgD-pgaA* region is a mutational hotspot across the Kp CC258 lineage

Given that PNAG production was widespread across Kp isolates, we next investigated the distribution and frequency of single nucleotide polymorphisms (SNPs) that have occurred across the *pgaABCD* operon, focusing on the high-risk CC258 lineage (Fig. 2A). To this end, we generated a mapping-based chromosomal alignment of 9214 CC258 genomes and used an ancestral reconstruction approach that, for each base position within the alignment, inferred the ancestral base state at all nodes across a recombination-free phylogenetic tree. We then analysed the occurrence of base (state) changes at each position across the Kp *pgaABCD* locus as well as the up and downstream regions, during the evolution and expansion of this lineage.

**Figure 2.**
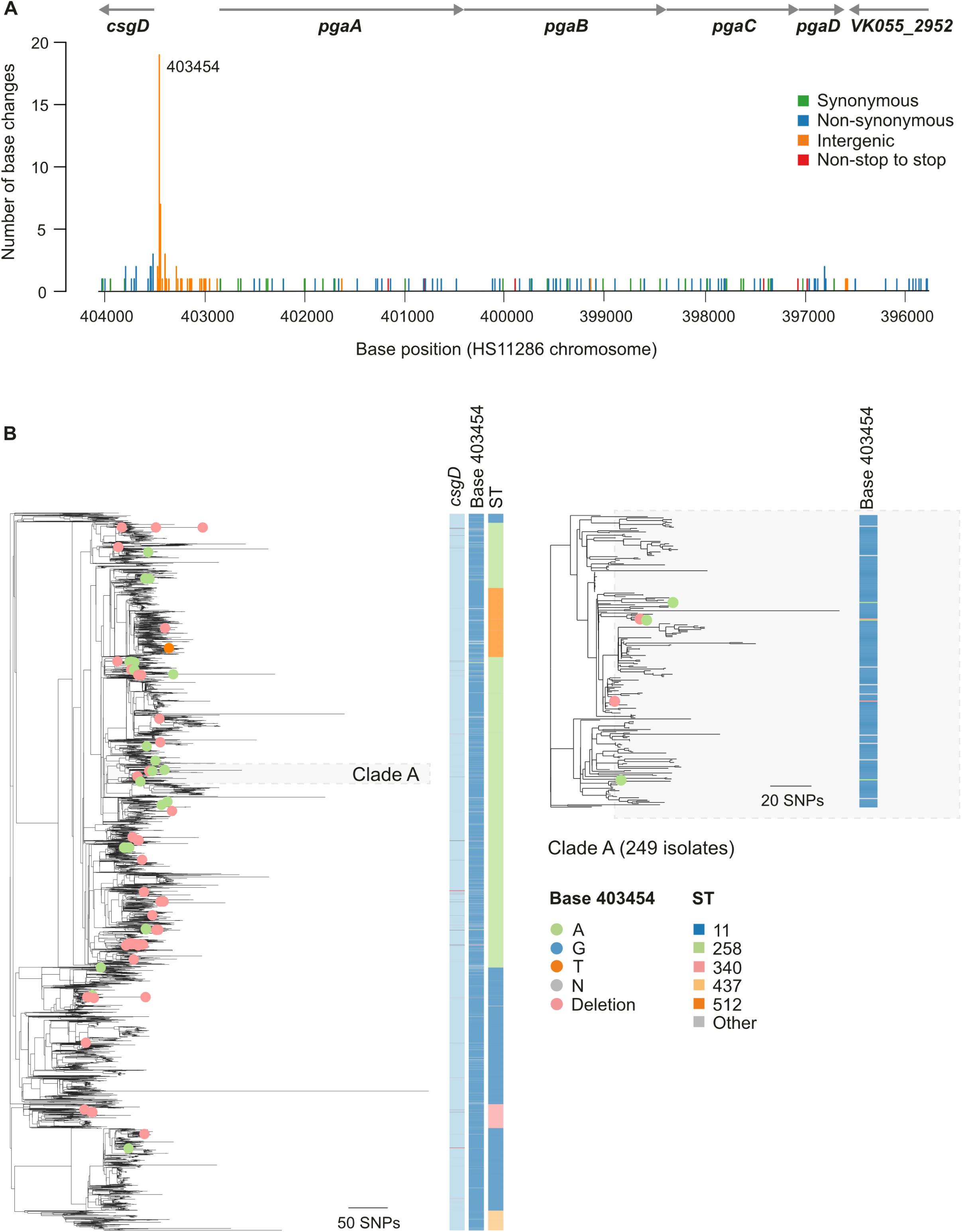
The intergenic *csgD-pgaA* region is a mutational hotspot across the Kp CC258 lineage. **(A)** Number of base (state) changes inferred to have occurred across the phylogeny of 9214 CC258 genomes at each base position spanning the length of the *pgaABCD* operon and up and downstream regions. Base changes are coloured by their effect on the protein sequence. **(B)** Phylogenetic tree of 9214 CC258 genomes constructed after the removal of recombined regions from the mapping alignment. Isolate nodes are coloured green if they possess an “A” or pink if they possess a deletion (-) at position 403454, with respect to the HS11286 reference. Metadata columns show the base called at position 403454, and the *csgD* gene (as characterised from the assemblies) and ST of the genomes. Clade A is shown with additional resolution on the right-hand side. The scale bars represent the number of SNPs. An interactive visualisation of this tree, together with additional metadata and genotypic data, is available via Microreact at https://microreact.org/project/kp-cc258-pnag.

We found that 98.48% of bases among the *pgaABCD* genes were conserved (Fig. 2A). In contrast, the IR spanning the start codon of *pgaA,* and the nearest upstream gene, a *luxR-*like transcriptional regulator annotated as *csgD*, underwent frequent mutation (Fig. 2A). In particular, we identified a position 36 bases upstream of *csgD* (position 403454 relative to the ST11 HS11286 reference), that has been subjected to 19 base changes across the CC258 phylogeny, indicative of a strong selection pressure acting on this position (Fig. 2A). Of the 19 base changes at this position, 17 constituted a G>A substitution, while there was a single A>G reversion, and one G>T substitution. Isolates encoding a G>A substitution occurred exclusively on or near terminal branches of the tree. We also observed that 44 genomes possessed a missing base at position 403454. This was typically associated with wider deletion events and the insertion of an IS element, resulting in partial or complete loss of *csgD*. Indeed, isolates possessing a missing base at this position were found to lack an intact *csgD* gene. Other nearby positions have also been subjected to multiple base changes including 403442 (position -48 of *csgD*) (7 changes) and 403451 (position -39 of *csgD*) (5 changes) (Fig. 2A and B). We also analysed the distribution of isolates carrying a G>A substitution at position 403454, or lacking *csgD* across the lineage. Isolates carrying either mutation emerged multiple times across the phylogeny, but interestingly, occurred exclusively on or near terminal branches of the tree. Therefore, isolates carrying a G>A substitution or lacking *csgD* failed to clonally expand, suggesting a medium- and/or long-term fitness cost and limit clonal expansion (Fig. 2B).

### Recurrent G>A mutation results in PNAG hyper-production

The genetic arrangement of *csgD* upstream of the *pgaABCD* structural genes resembles a typical bacterial operon structure, in which, the upstream transcriptional regulator modulates expression of the downstream genes. Therefore, we hypothesised that *csgD* was a transcriptional regulator of the *pgaABCD* genes, and that recurrent mutations in the region immediately upstream of *csgD*, impact PNAG production.

To test this hypothesis, we generated an isogenic ORF deletion in the *csgD* gene (Δ*csgD*) and introduced point mutations into ICC8001 to substitute the single base at position 403454 from a G to an A (-36_G>A_) or a T (-36_G>T_). These point mutations were used to model the frequent emergence of mutations that we observed at this position across CC258 genomes (Fig. 3A). Although the G>A substitution was most prevalent amongst ST258 strains, we engineered these mutants in a hvKp background to allow for subsequent virulence assessment of these SNPs, as this strain background provides a consistent and robust model of lethal infection (28).

**Figure 3.**
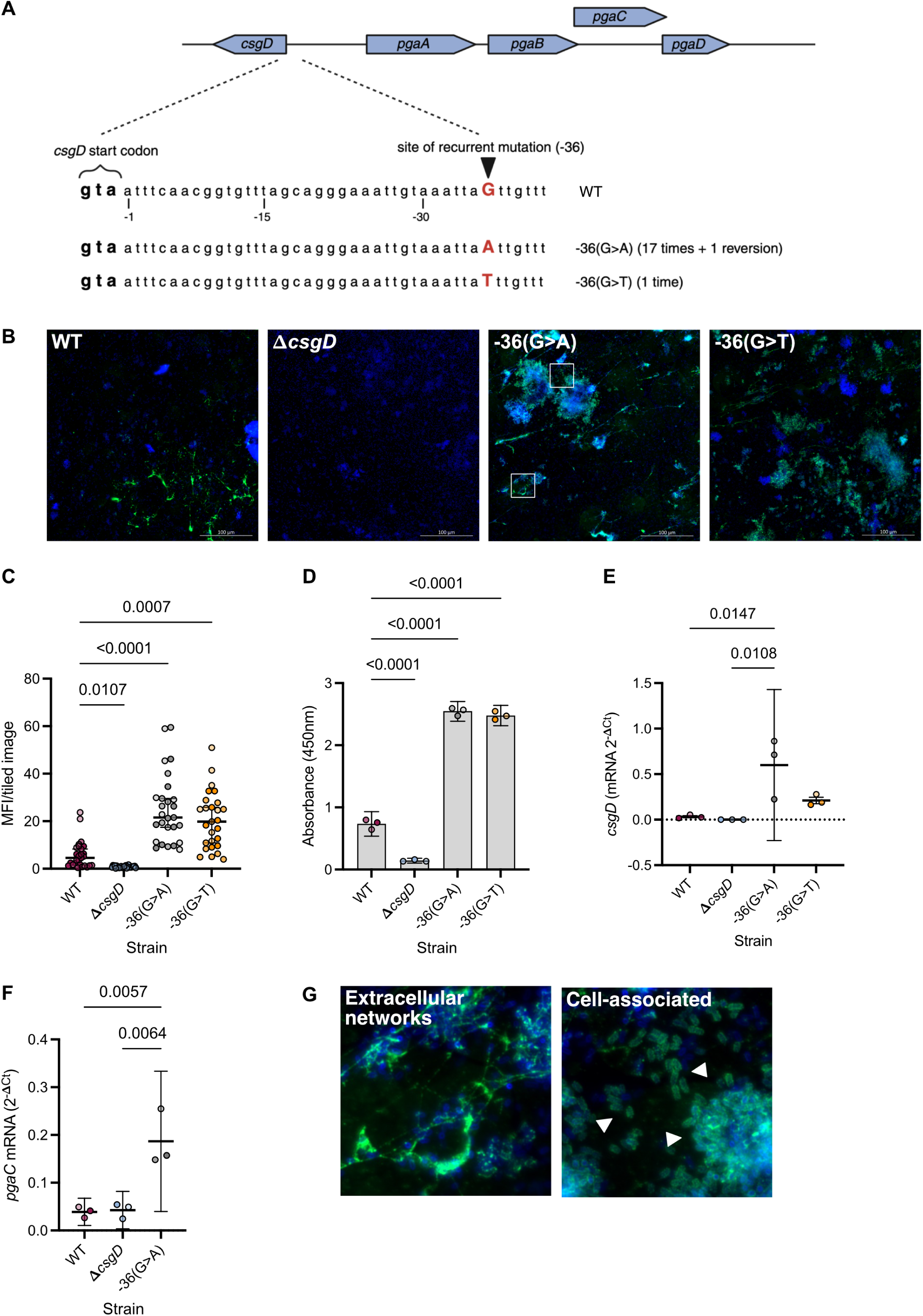
Recurrent G>A mutation upstream of *pgaR/csgD* results in PNAG-hyper-production. **(A)** Schematic of the Kp *pgaABCD* locus, including the upstream regulator *csgD,* and the diversity observed at position 403454 across CC258 genomes. **(B)** Individual tiled images of PNAG production in WT, Δ*csgD*, Kp -36_G>A_ and Kp -36_G>T_ after O/N static growth in LB on glass. Wells are stained with FM464 (blue) and F598 (green), and an individual tiled image, highlighted in Fig. S2, is shown. White squares highlight examples of extracellular PNAG networks and cell-associated PNAG, which are shown as zooms in **(G)**. **(C)** Quantification of PNAG fluorescence across wells. Individual points represent MFIs of each tiled image across 3 biological replicates. Points of the same shade are from the same replicate. Data is presented as median with 95% confidence intervals, and significance was assessed by Kruskal-Wallis. **(D)** ELISA absorbances of wells coated with Kp strains probed with 500 ng/mL mAb F598. Data is presented as mean with 95% confidence intervals across 3 biological replicates, which are shown in different shades. Significance was assessed using one-way ANOVA (p value <0.0001) with Tukey’s multiple comparisons. P values from post hoc testing are indicated on the graph. **(E)** *csgD* and **(F)** *pgaC* transcript levels, as measured by qRT-PCR in strains after O/N static growth on glass slides in LB. Data is presented as mean with 95% confidence intervals across 3 independent experiments. Significance was assessed using one-way ANOVA (p values 0.0089 and 0.0037 for *csgD* and *pgaC,* respectively), with Tukey’s multiple comparisons test. P values indicated on the graph are from Tukey’s post hoc tests. **(G)** Zoomed regions of **(B)** showing PNAG extracellular network formation and cell-associated PNAG production in Kp -36_G>A_-coated wells. White arrows highlight individual Kp producing cell-associated PNAG.

We grew the strains statically on glass slides and assessed PNAG production. While the WT strain produced PNAG extracellular networks, deletion of *csgD* abolished PNAG production and resulted in a strain that phenocopied our negative control (Δ*pgaBCD*) (Fig. 3B and S3). In contrast, IF imaging of Kp -36_G>A/T_ strains revealed widespread PNAG production across all well locations, at levels higher than WT-coated wells (Fig. 3B and S3). We also observed a change in growth phenotype in Kp -36_G>A/T_ coated wells, with regular formation of dense bacterial aggregates, consistent with improved intra-bacterial adherence (Fig. S3). Quantification of PNAG-specific fluorescence confirmed that loss of *csgD* abolished PNAG production, while Kp -36_G>A/T_ mutations resulted in significantly higher MFIs compared to WT (Fig. 3C). Similarly, we observed higher ELISA absorbances in wells coated with either Kp - 36_G>A_ or Kp -36_G>T_ when compared to WT-coated wells and negligible signal in Δ*csgD*-coated wells (Fig. 3D).

We also noted that the PNAG phenotype in Kp -36_G>A/T_ coated wells appeared different to WT-coated wells. Along with PNAG extracellular networks, we also observed more concentrated PNAG staining surrounding dense Kp aggregates (Fig. S3). Closer inspection of these regions revealed a cell-associated PNAG phenotype, which presented as circumferential staining of individual bacteria (Fig. 3G), which more closely mimics PNAG localisation on WT Kp in vivo (Fig. 1F) and on other bacteria (3,28). Considering that this phenotype is only visible on PNAG hyper-producing strains, we suggest that cell-associated PNAG production may be a product of elevated *pgaABCD* expression.

### *pgaR* (*csgD*) is a transcriptional regulator of the *pgaABCD* genes

Due to the proximity of G>A/T SNPs to the *csgD* start codon, we hypothesised that they may impact *csgD* expression, which could in-turn upregulate PNAG production. Indeed, we detected more *csgD* transcripts in Kp -36_G>A_ compared to WT (Fig. 3E). While *csgD* transcripts appear elevated in Kp -36_G>T_ cultures, they did not differ significantly from WT (Fig. 3E). Given that only the G>A substitution induced higher *csgD* expression, we also assessed expression of the PNAG biosynthetic gene *pgaC* in this genetic background. We detected higher *pgaC* transcripts in Kp -36_G>A_ compared to WT, consistent with increased PNAG expression (Fig. 3F). Therefore, the recurrent G>A mutation induces higher expression of *csgD,* and thus, PNAG biosynthetic genes, which results in a PNAG-hyper-producing phenotype.

Given that we did not detect PNAG on Kp Δ*csgD*, and that up-regulation of *csgD* results in increased *pgaABCD* expression, we concluded that the gene annotated as *csgD* in Kp is a novel positive regulator required for PNAG production. In *Escherichia coli* (*E. coli*) and *Salmonella enterica* serovar *Typhimurium* (*S.* Typhimurium), *csgD* is well characterised as a positive transcriptional regulator of the curli fimbriae synthesis and assembly genes (29,30). However, Kp lacks these genes and does not produce curli. On this basis we renamed *csgD* as *pgaR* (regulator of *pgaABCD*).

### -36_G>A_-induced PNAG hyper-production attenuates Kp virulence *in vivo*

As isolates lacking *pgaR* or carrying SNPs at position 403454 occur exclusively on or near terminal tree branches, we hypothesised that these strains may also have attenuated virulence. Therefore, we assessed how loss of PNAG (Δ*pgaBCD*) and PNAG-hyper-production (Kp - 36_G>A_) affect Kp virulence in *in vivo.* To test this hypothesis, we infected CD-1 mice with WT, Δ*pgaBCD* or Kp -36_G>A_ via IT intubation and assessed disease readouts 72 hpi. Infection with both WT and Δ*pgaBCD* induced similar weight loss, with means of 10.45% and 9.364%, respectively (Fig. 4A). In contrast, mice infected with Kp -36_G>A_ induced minimal weight loss, with a mean of 4.73% (Fig. 4A). Quantification of bacterial lung burden revealed no difference between WT and Δ*pgaBCD*, although Δ*pgaBCD* lung CFUs exhibited more variability (Fig. 4B). Notably, infection with Kp -36_G>A_ resulted in a significant reduction in lung CFU compared to WT, suggesting that over-production of PNAG attenuates Kp replication in the CD-1 lung (Fig. 4B). Furthermore, infection with WT and Δ*pgaBCD* consistently resulted in bacteremia, while only a single mouse infected with Kp -36_G>A_ became bacteremic (Fig. 4C). This equated to a significant reduction in blood CFUs following lung infection with Kp -36_G>A_ compared to WT (Fig. 4C).

**Figure 4.**
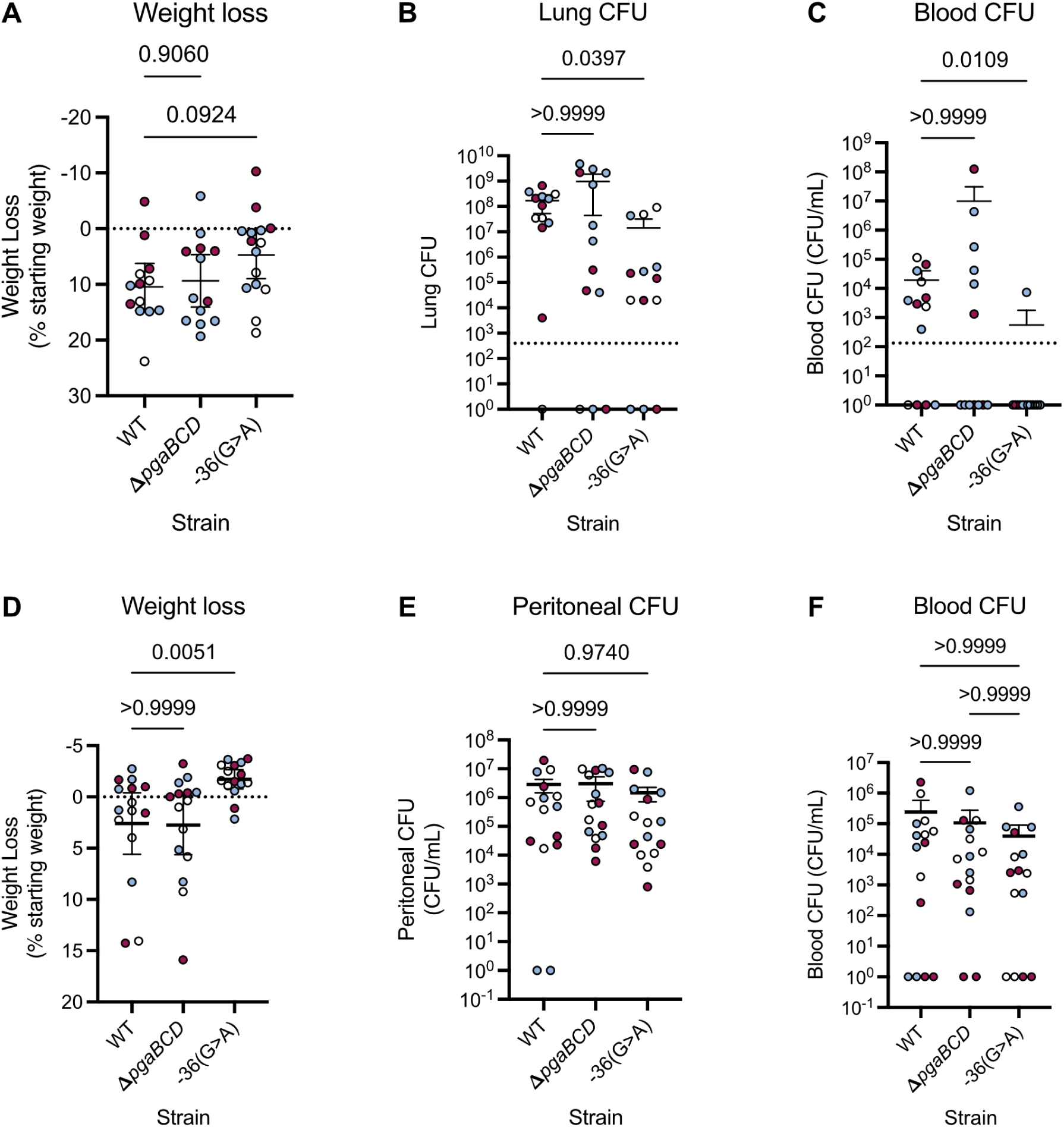
PNAG hyper-production attenuates Kp virulence *in vivo*. **(A)** Weight loss in CD-1 mice 72 hpi with 500 CFU Kp strains. Data is plotted as a percentage of mouse starting weight, with the mean and 95% confidence intervals shown. Significance was assessed using One-Way ANOVA (p value=0.1114) with Tukey’s multiple comparisons test. P values from post hoc tests are shown on the graph. Bacterial burden in the lungs **(B)** and blood **(C)** of CD-1 mice 72 hpi. Data are presented as individual data points with the mean and 95% confidence intervals indicated. Individual points are colour coded according to biological replicate. In the remaining cases, significance was assessed using non-parametric Kruskal-Wallis tests, with all p values are indicated on graphs.

We also assessed how these opposing PNAG phenotypes affected virulence during a murine peritonitis model. For this, we infected BALB/c mice via intraperitoneal (IP) inoculation, then quantified bacterial numbers in the peritoneum and blood 48 hpi. This revealed that mice infected with WT and Δ*pgaBCD* strains experienced significant weight loss, while Kp -36_G>A_ infected mice maintained their starting weight (Fig. 4D). Interestingly, infection with all three strains resulted in primary peritonitis and subsequent bacterial dissemination to the blood (Fig. 4E and F). Furthermore, bacterial burdens in both the peritoneum and bloodstream did not differ between strains (Fig. 4E and F). Therefore, while PNAG was dispensable for virulence in this setting, PNAG hyper-production alleviated the disease phenotype. Taken together, we find that PNAG is dispensable for Kp virulence during murine peritonitis and pneumonia models, but PNAG hyper-production attenuates Kp infection and the associated disease. This may help explain why the -36_G>A_ mutation does not expand in nature.

## Discussion

This study demonstrates that extracellular network formation is the predominant PNAG phenotype across Kp isolates. This contrasts with previous IF staining of bacterial PNAG, where PNAG presents as a cell-associated polysaccharide, often coating individual bacteria (3). This network phenotype also diNers markedly from the cellular localisation of other Kp surface polysaccharides like CPS and lipopolysaccharide (LPS), which circumferentially coat the bacterial surface and are associated with the outer membrane (28,31). Therefore, the extracellular network phenotype appears specific to PNAG production in Kp.

Across Gram-negative bacteria, the *pgaABCD* locus is subject to complex and multilayered regulation. In *E. coli*, the global carbon storage regulator CsrA binds to the 5′ untranslated region (UTR) of *pgaA* transcripts, blocking ribosome access and promoting transcript degradation, thereby repressing PNAG biosynthesis. In contrast, the LysR-type transcriptional regulator NhaR activates *pgaA* transcription in a Na⁺-dependent manner. Notably, both CsrA and NhaR are global regulators encoded at chromosomal loci distant from *pgaABCD* and control numerous additional targets. Here, we identify a positive transcriptional regulator of *pgaABCD*, designated *pgaR*, located immediately upstream of *pgaA* within the same operonic structure. This genomic organisation mirrors that of the *icaADBC* locus in Gram-positive bacteria such as *S. aureus*, where the divergently transcribed regulator *icaR* represses the structural genes required for PNAG synthesis. Importantly, whereas *icaR* functions as a local negative regulator, *pgaR* acts as a local positive regulator. Furthermore, *pgaR* appears to be restricted to the Kp species complex; no analogous upstream transcriptional regulator is present in reference strains of *E. coli*, in which PNAG production has been extensively characterised. This distinction highlights a lineage-specific regulatory adaptation in Kp that may enable more dynamic tuning of PNAG expression.

Across CC258 genomes, we identified 49 isolates either lacking *pgaR* or encoding a truncated variant. Given our experimental evidence that *pgaR* is required for PNAG production, and the broad phylogenetic distribution of these isolates, we infer that loss of PNAG has arisen independently multiple times within the CC258 lineage. Alongside these mutations, we observe recurrent mutations in the region upstream of *pgaR* that confers a PNAG hyper-producing phenotype. Therefore, across this lineage, we observe recurrent and opposing evolutionary trajectories: loss of PNAG through *pgaR* inactivation, and PNAG hyper-production driven by the Kp −36_G>A_ substitution upstream of *pgaR*. The limited clonal expansion of isolates carrying either truncated *pgaR* or the −36G>A mutation suggests that the selective advantage conferred by these phenotypes is transient or niche-specific, rather than broadly adaptive. We have previously reported that a recurrent synonymous mutation in the *ompK36* gene increases antibiotic resistance but attenuates Kp virulence and thus, fails to expand clonally (32). Consistent with this, we found that PNAG hyper-production attenuated Kp virulence and the associated disease during *in vivo* severe lung infection. This may explain the failure of isolates carrying a −36_G>A_ substitution to expand in nature.

Previous studies have reported mutations in regulatory regions associated with enhanced biofilm formation in Kp isolates from human urinary tract infections (UTIs), including alterations within the *pgaA–csgD* intergenic region (33). UTI-associated strains are typically characterised by increased biofilm-forming capacity, and in uropathogenic *E. coli* (UPEC), PNAG is an established virulence determinant (34). These observations suggest that the human urinary tract, or other infection niches in which surface attachment and biofilm maturation play a role, may favour a PNAG hyper-producing phenotype. Supporting this model, clinical isolates of *S. aureus* from cystic fibrosis (CF) patients have been found to harbour mutations in the *icaR–icaA* intergenic region, analogous to the *pgaR–pgaA* interval in Kp, that result in increased PNAG production (35,36). In contrast to the recurrent SNP observed upstream of *pgaR*, CF-associated *S. aureus* isolates typically carry a 5-bp deletion (TATTT motif) in the *icaA* promoter, abolishing binding of the negative regulator ROB (regulator of biofilm) and thereby relieving repression of *icaADBC* (35). Collectively, these findings indicate that mutations within regulatory regions upstream of PNAG biosynthetic loci represent a conserved evolutionary strategy to modulate biofilm formation across phylogenetically diverse, high-priority bacterial pathogens.

Regulatory evolution of biofilm-associated loci is not restricted to PNAG biosynthesis. Recurrent promoter mutations have also been described in the *csgD* promoter in other Gram-negative pathogens. For example, in *Salmonella enterica* serovar Typhimurium, a point mutation in the *csgD* promoter, identified as a -44_G>T_ substitution, increases *csgD* expression and produces a large, rough colony morphotype with enhanced biofilm-forming capacity. Notably, strains carrying a −44_G>T_ substitution in the *csgD* promoter have expanded into a high-risk lineage in China and continue to disseminate globally (37). In *S. Typhimurium*, CsgD functions as a master regulator with multiple downstream targets, coordinating curli production, cellulose synthesis, and broader biofilm architecture (38,39). The successful expansion of this lineage suggests that either the energetic cost of CsgD-regulated targets is lower than that of PNAG hyper-production in Kp, or that thFe ecological advantages conferred by enhanced CsgD activity outweigh associated fitness costs. In contrast, the absence of widespread expansion among Kp isolates harbouring the −36G>A mutation implies a substantial fitness trade-off associated with sustained PNAG overproduction.

Although we demonstrate that PNAG is dispensable during *in vivo* lung infection with the hvKp strain ICC8001, previous work has shown tFhat PNAG contributes to virulence in other Kp infection models, including peritonitis and gastrointestinal colonisation (25). In these contexts, loss of PNAG attenuates lethality in mice and reduces gut colonisation and subsequent systemic dissemination (25). Furthermore, the multiple virulence factors produced by our WT strainICC8001, including hypermucoidy and the siderophores yersiniabactin and salmochelin, have the potential to mask less prominent virulence determinants like PNAG (40). Together, these findings suggest that the contribution of PNAG to Kp pathogenesis is highly context dependent, influenced by factors like Kp strain background and the infection niche. While our *in vitro* observations showed that PNAG is forming extensive extracellular networks under static growth conditions, we found cell-associated PNAG following lung infection with WT ICC8001. Given that this phenotype was seen in during *in vitro* growth when PNAG was over-expressed, this suggests that the lung environment induces expression of PNAG in Kp.

Since PNAG production appears dispensable during acute lung infection, it may play a more prominent role in chronic or surface-associated infections. Therefore, infection models that better recapitulate persistent biofilm-associated disease, such as wound infection or device-associated colonisation, may therefore provide deeper insight into the conditions under which PNAG production enhances Kp fitness and virulence.

## Data availability

All the data are contained within the main and supplementary material. The ancestral state reconstruction across the tree can be found here: https:://github.com/sanger-pathogens/bact-gen-scripts/blob/master/reconstruct_snps_on_tree.py

## Acknowledgements

We thank Rita Berkachy for the excellent technical help. JB is supported by a BBSRC studentship, BB/W5101154/1. SD is funded by the Bill & Melinda Gates Foundation (investment number INV-025280). This study was supported by a grant from The Wellcome Trust 224282/Z/21/Z.

## Author contributions

JB – Conceptualisation, Data Curation, Formal Analysis, Investigation, Methodology, Writing – Original Draft Preparation

JSG – Data curation, Investigation, Writing – Review & Editing

SD – Data Curation, Formal Analysis, Methodology, Software, Investigation, Writing – review & Editing

RB – Investigation

MGP – Funding Acquisition, Supervision

IMR – Funding Acquisition, Writing – Review & Editing

MRR – Funding Acquisition, Supervision

JLCW – Supervision, Investigation, Conceptualisation, Writing – Review & Editing

GF – Supervision, Funding Acquisition, Conceptualisation, Writing – Review & Editing

## Methods

### Identification of homoplasic SNPs across the *pga* operon region among CC258 genomes

We assembled a collection of 9214 publicly-available CC258 genomes identified via Pathogenwatch (https://pathogen.watch) (41). We downloaded raw sequence reads associated with these genomes from the European Nucleotide Archive (ENA) using accession numbers available in Pathogenwatch. We also obtained the associated metadata and Kleborate v2.3.0 results from Pathogenwatch (42). The sequence reads were mapped to the ST11 reference genome, HS11286 (accession CP003200) using a workflow (https://gitlab.com/cgps/ghru/pipelines/snp_phylogeny) (version 1.2.2) incorporating BWA-MEM (43), SAMtools and BCFtools (44). We excluded isolates from downstream analyses if they had <20x average mapping depth or ≥25% missing sites in the resulting pseudo-genome alignment. We used Gubbins v3.2.1 (45) to remove recombined regions from the alignment and to generate a maximum likelihood phylogenetic tree with RAxML-NG (46). We subsequently used a maximum parsimony method to perform ancestral state reconstruction across the tree (https://github.com/sanger-pathogens/bact-gen-scripts/blob/master/reconstruct_snps_on_tree.py) for each base position within the alignment, thereby inferring the ancestral base (state) at each tree node. The number of state changes inferred to have occurred across the tree at each base position was calculated and the types of SNPs recorded. Base changes occurring within the region of the *pga* operon were then visualised and analysed.

### Bacterial growth conditions

All bacterial strains used in this study are listed in Table S1. Unless stated otherwise, all strains were cultured overnight (O/N) in Luria Bertani (LB) broth at 37℃, under agitation (200 rpm). When necessary, antibiotics were used at the following concentrations: streptomycin (50 µg/mL), gentamicin (10 µg/mL), rifampicin (50 mcg/mL).

### Kp mutant generation

All plasmids and primers used in this study are listed in Table S1. ORF deletion constructs were generated by assembling the 500 bp up and downstream of the gene of interest into linear pSEVA612SR, a pSEVA612S (JX560380.2) derivative with a flipped I-SceI site to reduce non-specific recombination, using Gibson Assembly (NEB). To generate point mutations in the *pgaR* upstream region, the *pgaR-pgaA* IR and flanking 500 bp up and downstream were assembled into linear pSEVA612SR. Site-directed mutagenesis, using primers listed in Table S1, was then employed to change the individual base at position 403454 from a G to an A or T. The resulting linear product was re-ligated using KLD reaction (NEB).

Gibson and KLD reactions were then transformed into chemically competent CC118λpir cells, which were then resuspended in 1 mL SOC media and recovered for 1 h at 37℃, shaking (200 rpm). Transformants were selected for by plating on LB agar supplemented with gentamicin (10 µg/mL). Mutagenesis plasmids were then introduced into ICC8001 using a two-step recombination protocol, as described previously (47). Successful recombinants were screened for size by colony polymerase chain reaction (PCR) and correct clones were confirmed by Sanger Sequencing (Eurofins).

### IF PNAG staining

Saturated Kp O/N cultures were diluted 1:1000 (v/v) into fresh LB and 200 µL was added to wells in a 96-well round glass bottom μ-plate (ibidi, #89607). Plates were incubated at 37℃ statically for 2 h, before media was aspirated and replaced with 200 µL fresh LB. Plates were then returned to the incubator O/N at 37℃, static. The next day, wells were aspirated and washed once in phosphate buffered saline (PBS) + 0.05% Tween-20 (PBST). Adherent Kp were then fixed in 4% PFA-PBS for 20 minutes (min) at room temperature (RT). Wells were washed three times in PBST and then probed with 100 µL mAb F598 (2 µg/mL), diluted in 1% Bovine Serum Albumin (BSA)-PBST O/N at 4℃. Wells were triple washed again in PBST, then incubated with 100 µL goat anti-human IgG AlexaFluor 488 (#A-11013, ThermoFisher Scientific), diluted 1:200 in 1% BSA-PBST, for 1 h at RT in the dark. Wells were washed three more times in PBST and incubated in FM464 (#F34653, ThermoFisher Scientific), diluted 1:100 in ultrapure water, for 20 min at RT in the dark. Wells were washed a final time in PBST and 200 µL sterile PBS added. Plates were imaged immediately using a Zeiss Axio Observer Z1 microscope.

For imaging, tiled images were taken at nine equally spaced, standardised fields across the well. MFIs of PNAG fluorescence across each field were calculated using the mean grey value function in FIJI (Version 1.54p) using the default thresholding parameters (48).

### Kp whole cell ELISA to detect PNAG

Kp O/N cultures were diluted and seeded into 96-well round glass bottom μ-plates (ibidi) as described above. After O/N static growth at 37℃, wells were washed once in PSBT and fixed in 4% PFA-PBS for 20 min at RT. Wells were triple washed in PBST then probed with mAb F598 (500 ng/mL) O/N at 4℃. The next day, wells were washed 3 times in PBST and incubated with goat anti-human IgG streptavidin HRP (#A18811, ThermoFisher Scientific), diluted 1:2000 into 1% PSA-PBST, for 1 h at RT, shaking (800 rpm). Next, wells were triple washed in PBST, and incubated with 3,3’,5,5’-Tetramethylbenzidine (TMB, #N301, ThermoFisher Scientific) at RT in the dark. Stop solution (50 µL) was added to terminate the reaction. Absorbance readings at 450 and 540 nm were taken immediately using a FLUOstar Omega plate reader.

### Quantitative real-time PCR (qRT-PCR)

Kp O/N cultures were diluted and grown statically on glass slides as described above. After O/N growth, adherent Kp were washed and scraped off slides, washed once in PBS, then resuspended in 500 µL PBS. Resuspended Kp were then treated with RNAProtect bacteria reagent (#76506, Qiagen) for 5 min at RT, then centrifuged at 10000 xg for 10 min. Bacterial pellets were then digested in 100 µL TE buffer with lysozyme (#L6876-1G, Sigma-Aldrich) (15 mg/mL) and 20 µL proteinase K (#19131, Qiagen) for 10 min with regular vortexing (every 2 min). RNA extraction was carried out using RNAeasy Qiagen Kit (#74104, Qiagen) according to the manufacturer’s instructions. Next, 1 µg RNA was treated with DNase (#M6101, Promega) for 1 h at 37℃ in the dark, and then cDNA synthesised using Moloney murine leukemia virus (MMLV) reverse transcription kit with random primers (#M1705, Promega). Real-time PCR was carried out using the Power Up SYBR Green master mix (#A25778, ThermoFisher Scientific) and primers specific for *pgaR, pgaC* and *Rho* as a housekeeping control (Table S1). The reaction was performed using a QuantStudio 1 System (Applied Biosystems) and results analysed using the QuantStudio Design & Analysis software (Applied Biosystems). Relative transcript levels were calculated using the 2^-ΔCt^ method.

### Animal experiments

All animal experiments were carried out in accordance with the Animals Scientific Procedures Act of 1986 (49) and UK Home Office guidelines (50) and were approved by the Imperial College Animal Welfare and Ethical review body (PP8896560). Specific pathogen-free female CD-1 (28-30g) were used for *in vivo* experiments and were housed in IVC cages with free access to food and water. Animals were housed with a 12h/12h light cycle, with temperature maintained at 20-24 C and humidity at 45-65%, in accordance with the UK Home Office code of practice. All animal experiments were reported in line with the ARRIVE guidelines (50). All inoculum for infection were prepared by diluting saturated Kp O/N cultures in sterile PBS to the desired concentration.

### *In vivo* severe lung infection

CD-1 mice were infected via IT intubation as described previously (28,32,51,52). Briefly, mice were anaesthetised via IP injection with ketamine (80 mg/kg) and medetomidine (0.8 mg/kg) and maintained at 32℃. Anaesthetised mice were suspended by their incisors and a canula was inserted into their trachea, with the use of a fibre optic cable. 50 µL diluted Kp O/N culture, containing 500 CFU, was pipetted into the canula and allowed to be spontaneously inhaled. Two 400 µL air flushes were then used to confirm complete inhalation. Anaesthesia was then reversed via subcutaneous injection of atipamezole (1 mg/kg), and mice were maintained at 32℃ until they had regained movement.

At 72 hpi, mice were placed under terminal anaesthesia via IP injection with ketamine (100 mg/kg) and medetomidine (1 mg/kg). Blood was collected from the heart via cardiac puncture and collected as described above. The thorax was opened and the lungs removed and collected into 3 mL sterile PBS in MACS C Tubes (Miltenyi Biotec). Lungs were homogenised, and along with blood samples, serially diluted and plated onto LB agar containing rifampicin (50 mcg/mL).

### *In vivo* intraperitoneal infection

Saturated Kp O/N cultures were diluted in sterile PBS to a final concentration of 250 CFU in 200 µL. BALB/c mice received 200 µL diluted WT, Δ*pgaBCD* or -36(G>A) Kp via IP injection. At 48 hpi, mice received anaesthesia via IP injection of ketamine (100 mg/kg) and medetomidine (1 mg/kg). Blood was collected directly from the heart via cardiac puncture, of which 20 µL was immediately mixed with 180 µL ultrapure water + 1 mM EDTA to enumerate CFUs. Finally, a 3 mL peritoneal lavage was performed, and 1 mL collected into Eppendorf tubes. Blood and peritoneal lavage samples were serially diluted and plated onto LB agar supplemented with rifampicin (50 mcg/mL) to quantify bacterial burden.

### Statistical analysis

All data analysis was performed using GraphPad Prism software (Version 5.10.0). All data is presented as mean ± sd unless otherwise stated. Prior to statistical analysis, all data was tested for normal distribution using a combination of D’Agostino-Pearson omnibus, Anderson-Darling, Shapiro-Wilk and Kolmogorov-Smirnov normality tests. Normally distributed data were analysed using parametric test, while non normally distributed data was analysed using non-parametric test. In each case, statistical tests used to assess significance are described in figure legends. Individual points are colour-coded to represent biological replicates.

## Supplementary figure legends

**Figure S1. Kp produces PNAG during static growth.** IF staining of PNAG in WT (A) and Δ*pgaBCD* Kp (B) at nine fields evenly distributed across wells, after static growth on glass slides in LB. Wells were stained with FM464 (blue) and mAb F598 (green). Images highlighted in red are those shown in Figure 1A. A single representative tiled well is shown from 3 independent experiments.

**Figure S2. Kp produces PNAG during an *in vivo* model of severe lung infection.** IF staining of lung sections of CD1 mice 72 h post IT intubation with 500 CFU Kp WT (A) and Δ*pgaBCD* Kp (B). Sections were stained with DAPI, podoplanin, Kp antiserum and mAb F598. Individual channels are shown in black and white, alongside a coloured merge with channel titles representing their respective colours in the image. A single representative image of lung infected with each strain is shown.

**Figure S3. Recurrent mutations in the *pgaR* promoter increase Kp PNAG production.** IF staining of PNAG in WT (A), Δ*csgD* (B), -36(G>A) (C) and -36(G>T) (D) Kp after O/N static growth on glass slides in LB. Images from nine fields, distributed evenly across the well are shown. Wells are stained with FM464 (blue) and mAb F598 (green), and tiles highlighted in red are shown in Figure 3. In each case, a single representative tiled well is shown from 3 biological replicates.

